# IRE1α-XBP1s axis is required for BCG-induced trained immunity against viral infection by inhibiting AHR signaling

**DOI:** 10.1101/2025.08.21.671435

**Authors:** Jincheng Chen, Chaohu Pan, Linghua Zhou, Hui Xu, Siqi Li, Junyan He, Min Huang, Yunlan Yi, Fuxiang Wang, Guoliang Zhang

## Abstract

Although Bacille Calmette-Guérin (BCG) has been proved to be suitable in inducing trained immunity, mechanism-related issues have not yet been fully resolved. In this study, we found that in vitro BCG training protects both BCG harboring and bystander macrophages against DNA virus and RNA virus infection and demonstrated that BCG harboring cells protect bystanders partly by type I interferon (IFN). We determined aryl hydrocarbon receptor (AHR) as a negative regulator in BCG-induced trained immunity on defense against viral infection in vitro and in vivo and blockade of type I IFN signaling partially impaired the enhanced antiviral effect of AHR deficiency in BCG training. Furthermore, we revealed IRE1α-XBP1s signaling as a critical pathway in BCG-induced trained immunity against viral infection in vitro and in mice. Mechanistically, we demonstrated that IRE1α-XBP1s pathway influenced BCG-induced trained immunity by limiting AHR mRNA expression and the translocation of AHR protein into the nucleus.

## Introduction

Innate immune cells have been observed to present a phenomenon of immune memory similar to that of adaptive immune cells, referred to as ‘trained immunity’ [1]. Candida albicans infection induced function reprogramming of monocytes, which resulted in increased production of cytokine in vitro and in vivo and enhanced protection against Candida albicans reinfection [2].

Other substances besides Candida albicans have also been found to be capable of inducing trained immunity, such as bacterial mucosal vaccines [3] or BCG [4]. In murine models, intravenous injection of BCG was more effective in inducing trained immunity and protecting mice against the attack of influenza virus than subcutaneous injection and the control group. However, BCG-induced trained immunity did not provide good protection for mice against the attack of the Severe Acute Respiratory Syndrome Coronavirus-2 (SARS-CoV-2) [5]. Nevertheless, some have reported that BCG-induced trained immunity through intravenous injection could effectively protect wild-type mice or ACE2 humanized mice against the attack of SARS-CoV-2, significantly reducing the disease score, decreasing the infiltration of inflammatory cells in the lungs, reducing weight loss, and enhancing the survival rate [6]. In a randomized placebo-controlled experiment, the monocytes of individuals vaccinated with BCG were found to undergo extensive epigenetic reprogramming and protect the subjects against experimental attenuated yellow fever virus infection in humans. The reduction of viremia was highly correlated with the upregulation of IL-1β rather than IFN-γ. The authors also demonstrated from the aspects of genes, epigenetics, and immunology that IL-1β is an important factor in inducing trained immunity [7]. NOD2 is the receptor for monocytes and macrophages to recognize BCG and induce trained immunity. Blocking TLR2, TLR4, or knocking out Dectin-1 could not eliminate the ability of BCG to induce trained immunity, but monocytes with NOD2 functional defects due to frameshift mutations failed to trigger a stronger cytokine response (i.e., impaired trained immunity function) after BCG treatment [8]. These findings suggested that BCG-induced trained immunity could provide protection against pathogens infection. However, the mechanism has not been fully investigated.

Here we explored the role of BCG-induced trained immunity in defense against viral infection. Employing mCherry-expressing BCG, GFP-expressing HSV-1 and GFP-expressing vesicular stomatitis virus (VSV), we found that BCG training could protect macrophages from DNA and RNA virus infection without requiring BCG living inside the cells. Furthermore, bystander cells were protected by BCG-harboring cells partly via type I IFN signaling in BCG-induced trained immunity upon viral infection. We observed that AHR deficiency enhanced the effect of trained immunity induced by BCG in vitro and in mice against viral infection. The IRE1α-XBP1s signaling is the most conserved pathway in the unfolded protein response (UPR) and reported to be important in immune response, however, its role in trained immunity remains unclear. Here we found that IRE1α-XBP1s signaling is critical in BCG-induced trained immunity against viral infection in vitro and in vivo. Mechanistically, IRE1α-XBP1s signaling inhibits AHR mRNA expression and AHR protein translocating into the nucleus in BCG-induced trained immunity, leading to elevated type I IFN production which limits viral infection.

## Results

### BCG training protects macrophages against DNA and RNA virus infection

To model trained immunity in vitro, human monocytic cell line Tohoku Hospital Pediatrics-1 (THP-1) cell-differential macrophages were exposed with 1 or 5 multiplicity of infection (MOI) mCherry-expressing BCG for 24 hours, after resting for 5 days, the cells were infected with GFP-expressing HSV-1 virus, a DNA virus (Fig. 1a and b). We observed that THP-1 treated with 5 MOI BCG showed significantly lower GFP positive percentage of cells and reduced viral titers 24 hours and 48 hours after HSV-1 infection (Fig. 1c). These indicated that suitable BCG training could induce some kind of ‘memory’ of macrophages for coping with subsequent viral attack. So, 5 MOI BCG was chosen as standard training dosage for subsequent experiments. Consistent with the above results, BCG training reduces GFP positive cells as demonstrated by immunofluorescence imaging (Fig. 1d). THP-1 trained with BCG also showed decreased viral infection rate and viral titers upon RNA virus (GFP-expressing VSV) infection (Fig. 1e) and these results were further confirmed by immunofluorescence imaging (Fig. 1f). The effect of trained immunity induced by BCG against viral infection was also confirmed on two other monocytic cell lines, U937 and RAW264.7, but not on nonimmune cells, A549 (Fig. 1g), although BCG could also be detected in A549 cells with flow cytometry 5 days after BCG exposure. To determine whether trained immunity induced by BCG treating against viral infection depends on direct BCG infection, we treated the THP-1 cells with rifampicin 24 hours after BCG exposure to kill the organism. The result showed that direct BCG infection is not required for BCG training-mediated protection (Fig. 1h).

**Fig.1.**
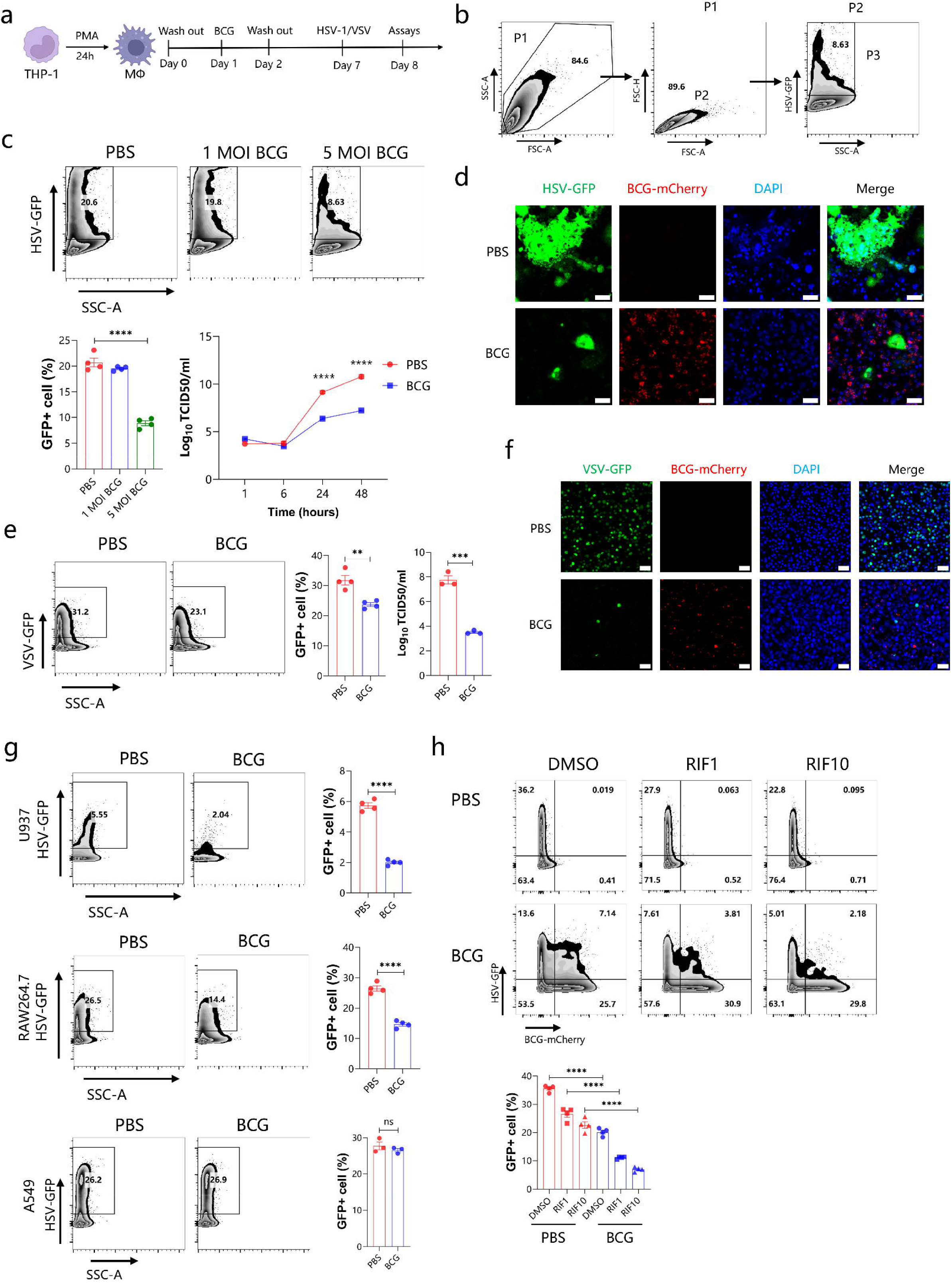
BCG training protects macrophages against DNA or RNA virus infection. THP-1-differentiated macrophages were trained with 1 or 5 MOI mCherry-expressing BCG for 24 hours, after 5 days resting, the cells were infected with 5 MOI GFP-expressing HSV-1 and harvested for assays (a). Gating strategy for analysis of HSV-GFP positive cells (b). The percentage of HSV-GFP positive cells were determined by flow cytometry analysis and viral titers were assayed by TCID_50_ (c). Immunofluorescence imaging to visualize BCG or HSV-1 infected cells (d). The percentage of VSV infected cells and VSV titers were determined (e). Immunofluorescence imaging to visualize BCG or VSV infected cells (f). The percentage of HSV-1 infected cells were checked in U937, RAW264.7, A549 cells (g). THP-1 cells were treated with 1 or 10 μg/ml Rifampicin (RIF1, RIF10) after BCG training, after 5 days resting, cells were infected with HSV-1 and checked with cytometry to determine viral infection rate (h). Scale bar: 50μm. Data are representative of three independent experiments and showed as mean ± SEM. *p < 0.05, **p < 0.01, ***p < 0.001, ****p < 0.0001 by *student t* test or one-way ANOVA.

We also performed RNA sequencing for characterizing transcription profile of BCG trained THP-1 upon HSV-1 infection and found that the transcriptome profile of BCG-trained THP-1 cells differed from the control group (Fig. S1a and b). Gene Ontology (GO) Cellular Component analysis revealed that BCG training was associated with nucleoplasm, cytosol, nucleus and cytoplasm (Fig. S1c). Furthermore, Kyoto Encyclopedia of Genes and Genomes (KEGG) analysis showed that BCG-trained THP-1 cells had stronger signal transduction, infectious disease (viral), immune response and transport catabolism related genes expression than the control (Fig. S1d, e and f).

These findings suggested that macrophages, but not nonimmune cells trained with BCG could induce trained immunity against DNA and RNA viral attack independent of BCG living inside the cells and BCG training resulted in differential transcription characteristics.

### BCG training protects both bystander cells and BCG harboring cells against viruses with a distinct profile

With mCherry-expressing BCG, we analyzed the different antiviral capacity between BCG-infected cells and non-infected bystander cells. The results revealed that both bystanders and BCG-infected cells in BCG training group exhibited decreased HSV-1 infection compared to the control (Fig. 2a). Similar results were also observed in VSV infection (Fig. 2b). Interestingly, we found that the percentage of HSV-1-GFP positive cells in BCG-infected cells was higher than that in bystander cells upon HSV-1 infection (Fig. 2a), which was contrary to the result upon VSV infection (Fig. 2b). While, we observed that the percentage of phosphorylation of Interferon Regulatory Factor 3 (p-IRF3) and IFN-β positive cells in BCG infected cells were increased compared to bystanders upon HSV-1 (Fig.2c and Fig. S2a) and VSV infection (Fig. 2d and Fig. S2b), which suggested that the reason why BCG-infected cells posed higher viral infection rate in HSV-1 infection might not be due to decreased immune response. Then we speculated BCG infection impacting viral entry receptors expression. In line with our speculation, the expression of HSV-1 entry receptors of PILRA and TNFRSF14 were enhanced in BCG-trained group than the control upon HSV-1 infection, while the VSV entry receptor of LDLR had no significant difference upon VSV infection compared to the control (Fig. 2e and Fig. S2c). Furthermore, we found that the percentage of PILRA and TNFRSF14 positive cells were higher in BCG-infected cells than bystanders upon HSV-1 infection, but that of LDLR had no significant difference upon VSV infection (Fig. S2d). The results were further confirmed by relevant siRNA experiment (Fig. 2f and g).

**Fig. 2.**
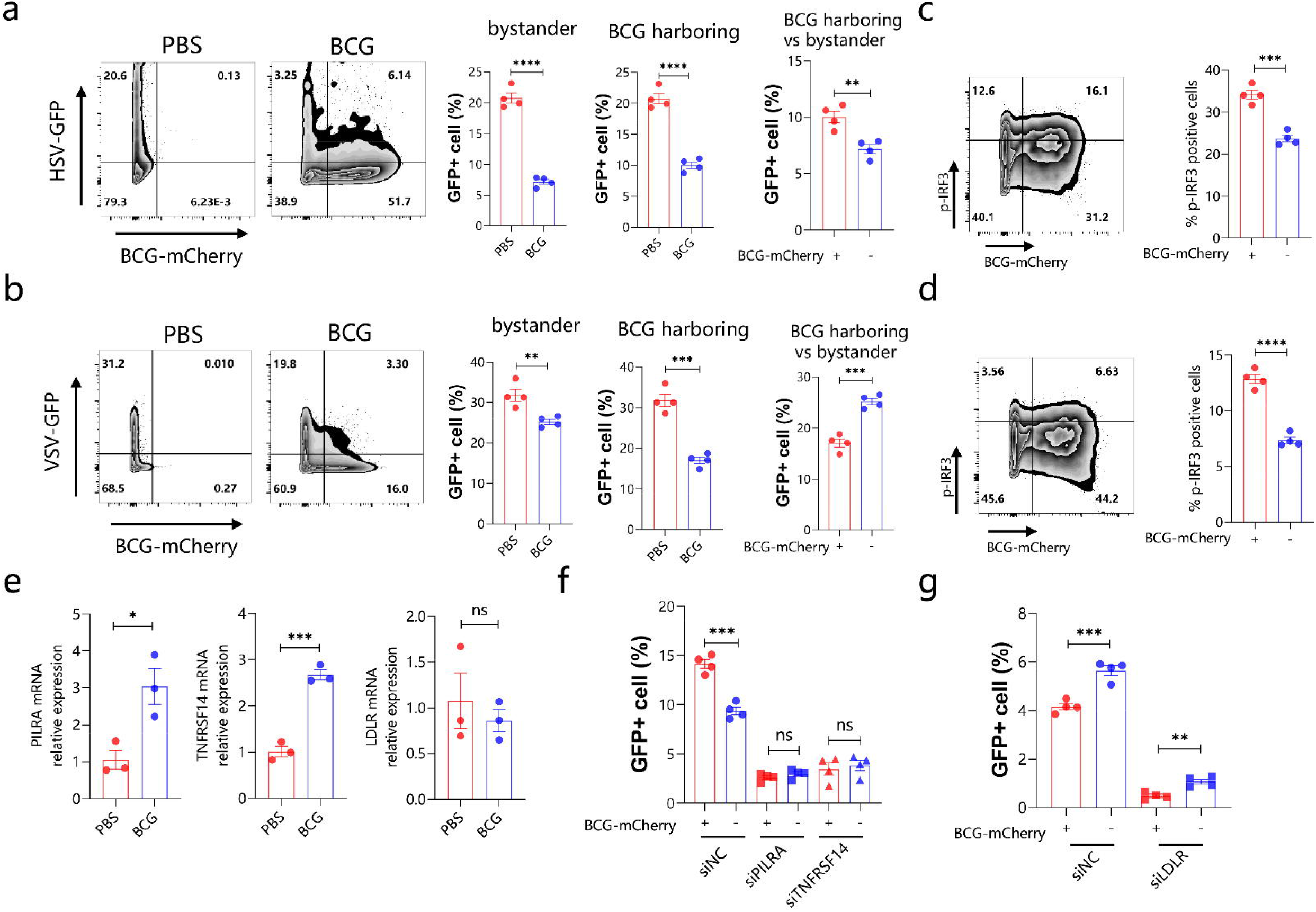
BCG training protects both bystander cells and BCG harboring cells against virus infection with a distinct profile. THP-1-differentiated macrophages were trained with BCG and rested for 5 days and infected with HSV-1 (a) or VSV (b), the percentage of viral infected cells in BCG-mCherry negative cells (bystander), BCG-mCherry harboring cells were investigated. 8 hours after HSV-1 (c) or VSV (d) infection, the percentage of p-IRF3 positive cells between bystander and BCG-mCherry harboring cells were compared. The mRNA expression of PILRA, TNFRSF14 and LDLR genes in BCG training cells and control were measured by qPCR (e). Cells trained with BCG were transfected with siRNA against PILRA, TNFRSF14 or negative control and rested for 5 days and infected with HSV-1, then the percentage of viral infected cells between bystander and BCG-mCherry harboring cells were compared (f). Cells trained with BCG were transfected with siRNA against LDLR or negative control and rested for 5 days and infected with VSV, then the percentage of viral infected cells between bystander and BCG-mCherry harboring cells were compared (g). Data are representative of three independent experiments and showed as mean ± SEM. *p < 0.05, **p < 0.01, ***p < 0.001, ****p < 0.0001 by *student t* test.

Then we attempted to study the capacity of bystander cells against viral infection derived from itself establishing trained immunity (innate immunity memory) or from BCG-infected cells. We used a setup in which donor cells were infected with BCG for 24 hours, and the supernatants were subsequently harvested and filtered with 0.22μm to prevent BCG and transferred to recipient cells for 24 hours, then the recipient cells were rest for 5 days and infected with HSV-1. The supernatants from donor cells trained with BCG could not provide significant protection for recipient cells against HSV-1 infection in THP-1, U937 and RAW264.7 cells and VSV infection in THP-1 cells (Fig. S3 a-e), suggesting that the bystander cells might not acquire immune memory in BCG training.

These results revealed that BCG training on the one hand could protect both BCG-infected and bystander cells against DNA and RNA viral infection, on the other hand influence alteration expression of some viral entry receptors, which results in distinct profile of viral infection between BCG-infected cells and bystanders.

### BCG harboring cells protect bystanders partly by type I IFN

Given that type I IFN signaling plays critical role on against viral infection, we evaluated the activated profile of the type I IFN in BCG-induced trained immunity. The results showed that BCG-trained cells exhibited higher IFN-β level 24 hours after HSV-1 infection and enhanced interferon-stimulated genes (OAS1, IFIT1) mRNA expression 6, 12 and 24 hours post-HSV-1 infection (Fig. 3a and b). Given the observation that bystander cells do not acquire trained immunity (Fig. S3 a-e), we sought to determine how bystander cells were protected. We added cells stained with CellTrace (as bystander cells) into the cells trained with BCG or control group during HSV-1 infection and checked the percentage of viral infected cells (GFP positive cells) of total CellTrace-positive cells by flow cytometry (Fig. 3c and d). Wildtype (WT) THP-1 or A549 CellTrace-stained cells transferred into BCG-trained THP-1 cells posed decreased viral infection than the control group (Fig. 3e and f), while blocking of type I IFN signaling (IFNAR1 knockout (KO)) markedly impaired this protection (Fig. 3e and f). Together, our results showed that bystander cells acquired protection from BCG-trained cells against viral infection and this process is partly dependent on type I IFN signaling.

**Fig. 3.**
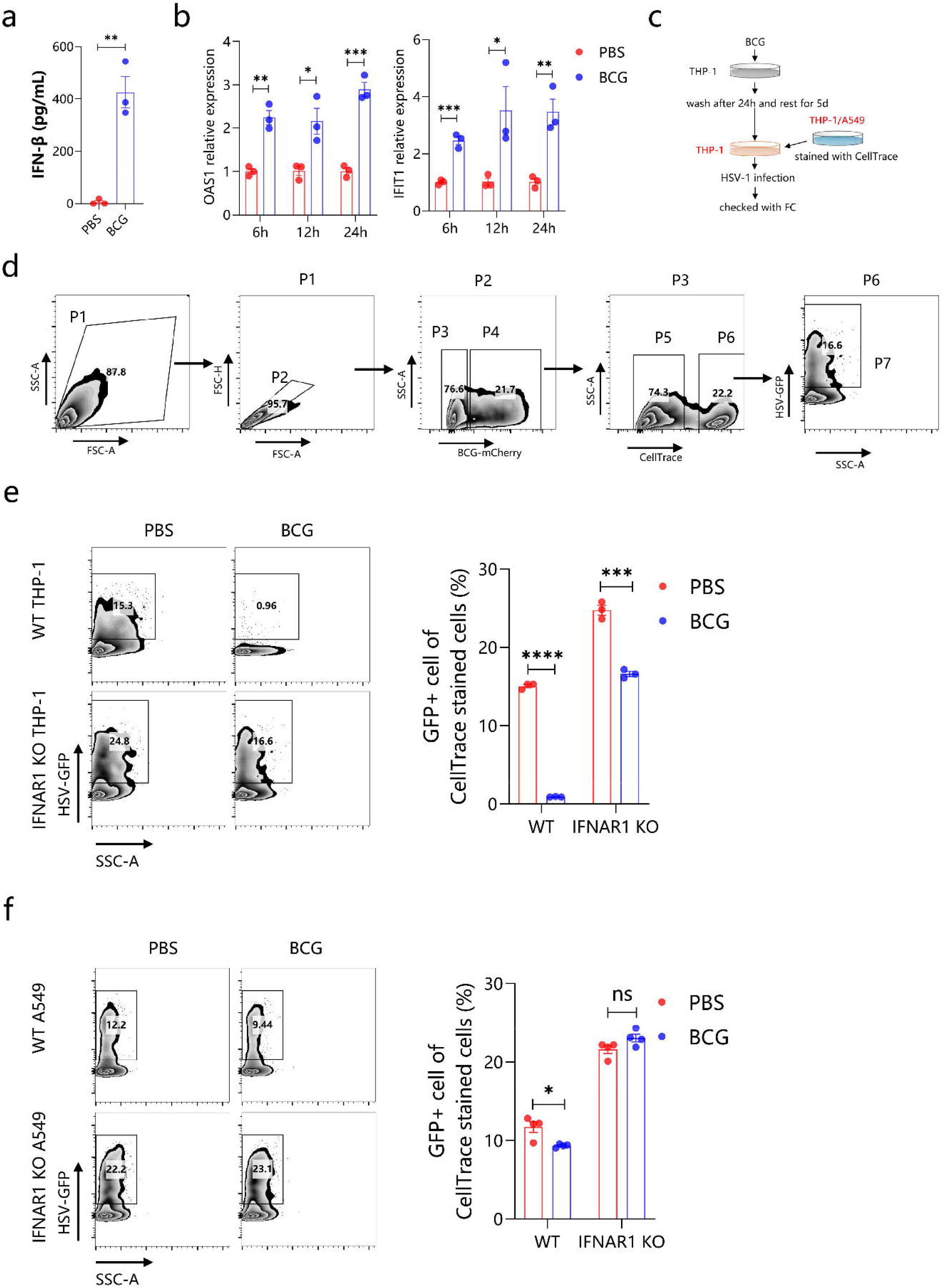
BCG harboring cells protect bystanders partly by type I IFN. IFN-β protein level (a) or ISGs OAS1 and IFIT1 mRNA expression (b) were assayed. THP-1 or A549 cells were stained with CellTrace and transferred into BCG trained THP-1 cells or control group and then infected with HSV-1 immediately, the viral load of CellTrace positive cells were measured by cytometry (c). Gating strategy for analysis of viral load of CellTrace positive cells (d). The viral load of CellTrace positive cells in THP-1 cells (e) or A549 cells (f) experimental systems. Data are representative of three independent experiments and showed as mean ± SEM. *p < 0.05, **p < 0.01, ***p < 0.001, ****p < 0.0001 by *student t* test.

### Deficiency of aryl hydrocarbon receptor (AHR) enhanced trained immunity against viral infection partly via type I interferon

In our previous study, we had observed that blockade of AHR signaling led to reduced viral infection, which indicated that AHR pathway could negatively regulate antiviral immunity [9]. Thus, we sought to evaluate the role of AHR signaling on BCG-induced trained immunity against viruses in vitro. Pharmacological inhibition or genetic knockout of AHR synergistically reduced viral loads in HSV-1 (Fig. 4a) or VSV infection (Fig. 4b) compared to BCG training alone group. Furthermore, AHR KO THP-1 cells trained with BCG led to enhanced IFN-β level upon HSV-1 infection (Fig. 4c) and blockade of type I IFN signaling with specific antibody during HSV-1 infection resulted in increased viral infection rate not only in wild type (WT) cells but also in AHR KO cell trained with BCG (Fig. 4d). These results revealed that the synergistically protective effect of AHR deficiency on BCG-induced trained immunity is partly dependent on type I IFN signaling.

**Fig. 4.**
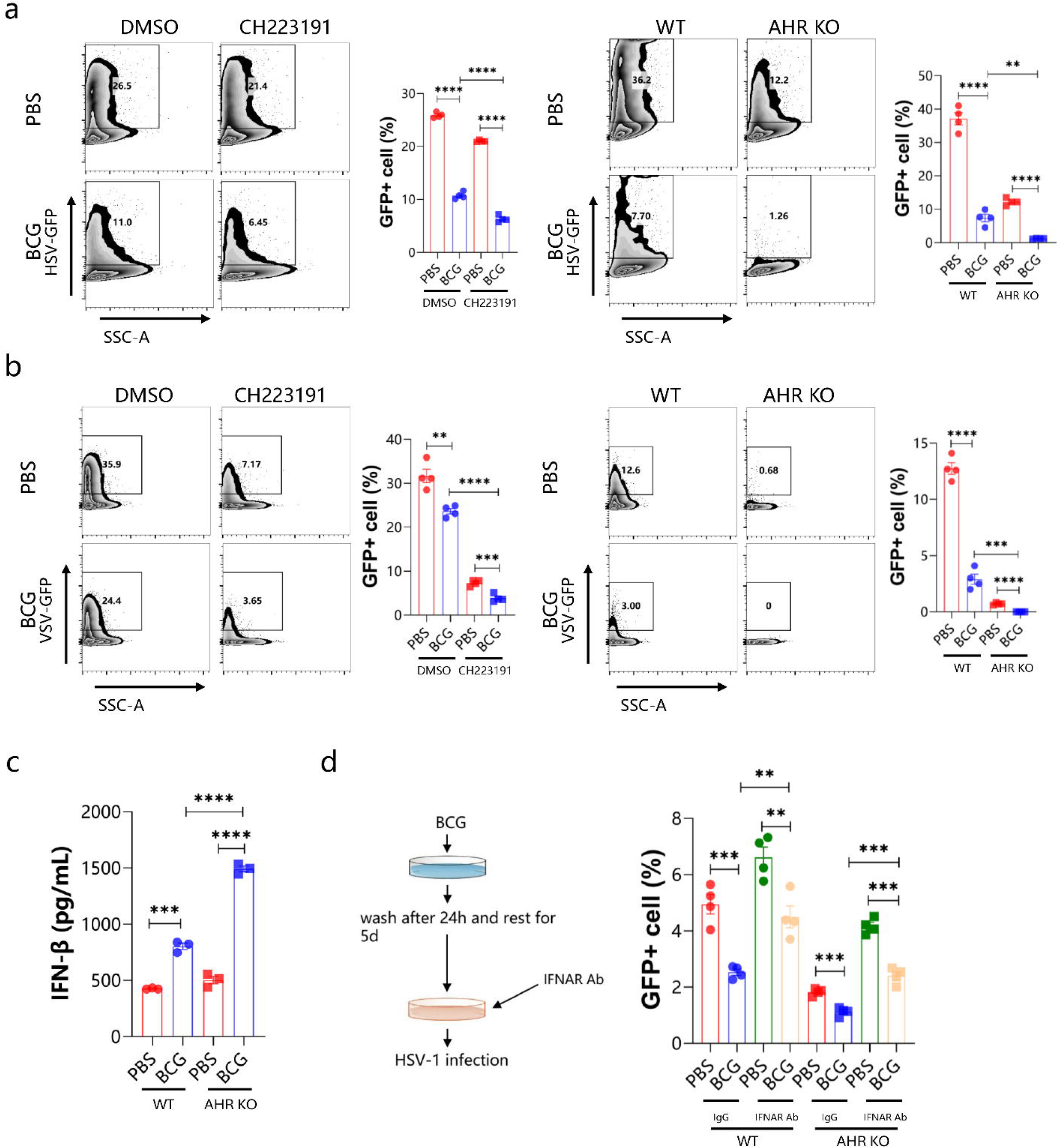
Deficiency of aryl hydrocarbon receptor (AHR) enhanced trained immunity against viral infection partly via type I interferon. THP-1-differentiated macrophages pre-treated with CH223191 (AHR antagonist) were trained with BCG for 24 hours and rested for 5 days, followed by HSV-1 infection (left); WT or AHR KO THP-1-differentiated macrophages were trained with BCG for 24 hours and rested for 5 days, followed by HSV-1 infection (right) (a). THP-1-differentiated macrophages pre-treated with CH223191 (AHR antagonist) were trained with BCG for 24 hours and rested for 5 days, followed by VSV infection (left); WT or AHR KO THP-1-differentiated macrophages were trained with BCG for 24 hours and rested for 5 days, followed by VSV infection (right) (b). The level of IFN-β was determined by ELISA in BCG training WT or AHR KO cells or control upon HSV-1 infection for 24 hours (c). Type I IFN signaling was blocked by using anti-IFNAR antibody in HSV-1 infection, and viral load was assayed by cytometry (d). Data are representative of three independent experiments and showed as mean ± SEM. *p < 0.05, **p < 0.01, ***p < 0.001, ****p < 0.0001 by one-way ANOVA.

To further confirm the effect of AHR signaling on BCG-induced trained immunity, we established AHR ^flox/flox^ and Lyz2 Cre AHR ^flox/flox^ mice and treated these mice with a single dose of BCG intravenously (Fig. 5a). These mice were challenged with HSV-1 virus after 35 days. In line with the in vitro data, mice whose macrophages deficient of AHR (Lyz2 Cre AHR ^flox/flox^) trained with BCG showed decreased percentage of HSV-1 infected cells (Fig. 5b), reduced relative viral DNA copy (Fig. 5c), and lower viral titer (Fig. 5d) than the WT control treated with BCG. We further observed that Lyz2 Cre AHR ^flox/flox^ mice trained with BCG exhibited higher IFN-β level 3 days after HSV-1 infection (Fig. 5e). Mice trained with BCG showed milder lung damage determined by H&E staining upon HSV-1 infection (Fig. 5f).

**Fig. 5.**
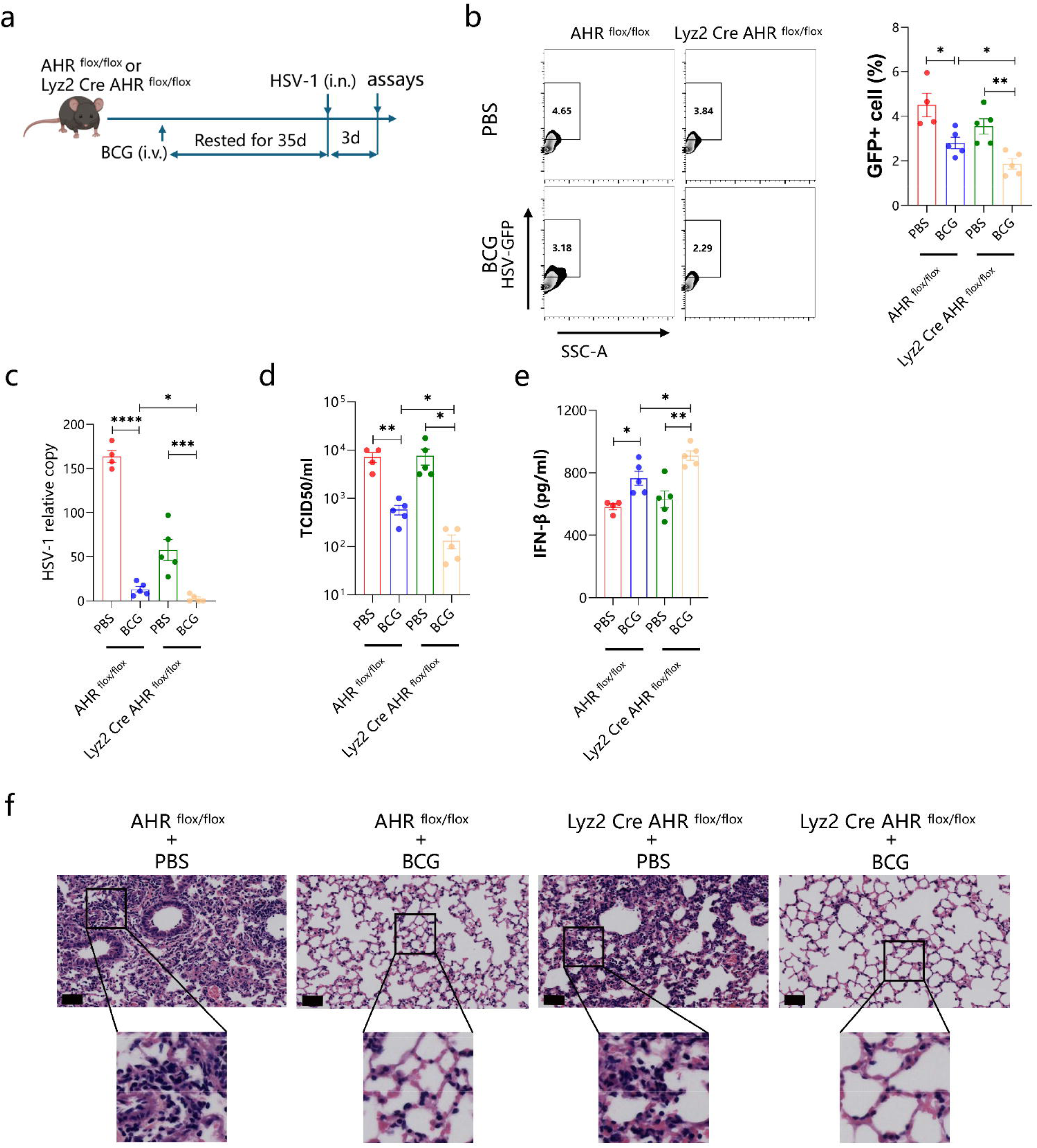
AHR deficiency on macrophages enhanced BCG-induced trained immunity against viral infection. 4-5 AHR ^flox/flox^ or Lyz2 Cre AHR ^flox/flox^ mice per group were trained with BCG or PBS as control intravenously. Rested for 35 days after BCG training, mice were intranasal infected with HSV-1 for 3 days and lungs were harvested for further assays (a): flow cytometry analysis (b), viral DNA copy assay by quantitative PCR for gD gene(c), viral titers by TCID_50_ assay (d), IFN-β level by ELISA (e), and pulmonary pathology by H&E staining (f). Scale bar: 50μm. Data are representative of three independent experiments and showed as mean ± SEM. *p < 0.05, **p < 0.01, ***p < 0.001, ****p < 0.0001 by one-way ANOVA.

As reported that histone 3 lysine 4 trimethylation (H3K4me3) and histone 3 lysine 27 acetylation (H3K27ac) were two primary epigenetic modifications in BCG-induced trained immunity, which facilitated chromatin accessibility and promoted specific genes expression [7, 8, 10, 11]. In our study we found that inhibition of H3K4me3 with MTA or H3K27ac with SGC-CBP30 could not significantly influence viral infection and IFN-β level not only in WT cells but in AHR KO cells trained with BCG (Fig. S4a and b).

### IRE1α-XBP1s signaling affects BCG induced trained immunity by AHR signaling

To elucidate mechanism regulating AHR signaling in BCG training, pathways involved in gap junctions, ion transport, endoplasmic reticulum stress, exosome biogenesis/release were checked in our study (data not showed), we found that activated the IRE1α-XBP1s pathway with APY29, one of the pathway in UPR, could significantly reduce viral infection in BCG-trained WT cells but not in BCG-trained AHR-KO cells in vitro. We confirmed this result with a selective antagonist for IRE1α-XBP1s pathway (4μ8C) and observed that inhibition of IRE1α-XBP1s pathway was able to profoundly increase viral infection in BCG-trained WT cells but only slightly increase in BCG-trained AHR-KO cells (Fig. 6a), suggesting that IRE1α-XBP1s pathway was critical in BCG-induced trained immunity by regulating AHR. We further employed three pharmacological inhibitors to target different arms of UPR. Blockade of IRE1α-XBP1s signaling with 4μ8C or ATF6 signaling with CaepinA7 during BCG training led to dose-dependent increase in viral infection rate (Fig. 6b). The results were confirmed with another IRE1α-XBP1s signaling inhibitor AMG18 and siRNA that target the XBP1 mRNA (Fig. 6c and d). In light of these findings, we focused on the IRE1α-XBP1s pathway. we further found that interruption of IRE1α-XBP1s signaling with 4μ8C leaded to decreased IFN-β level upon HSV-1 infection (Fig. 6e). We also observed that antagonizing IRE1α-XBP1s pathway during BCG training caused increased VSV infection rate (Fig. 6f).

**Fig. 6.**
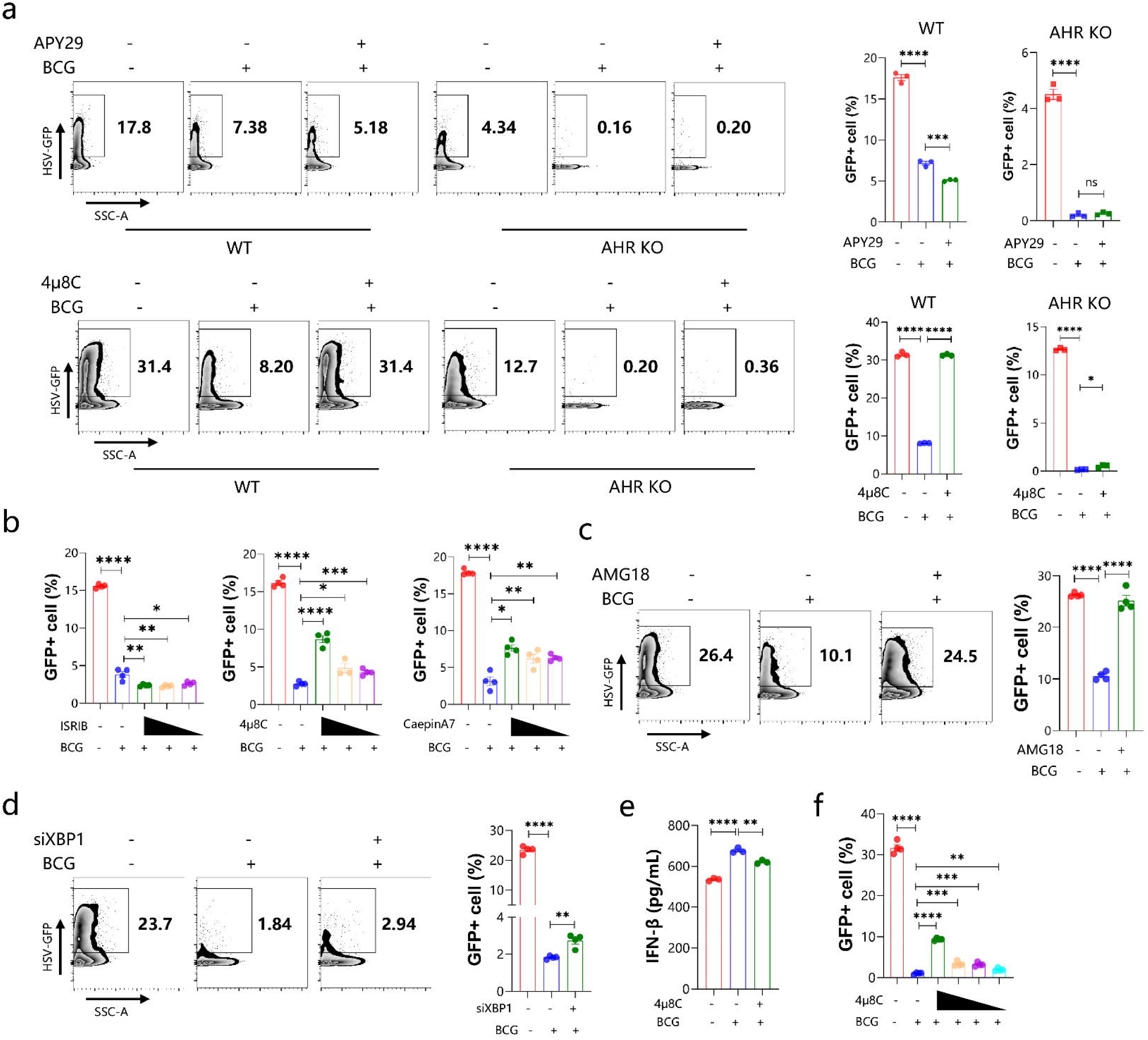
IRE1α-XBP1s signaling affects BCG induced trained immunity by AHR signaling. WT or AHR KO THP-1 differentiated macrophages pre-treated with APY29 or 4μ8C were trained with BCG, then rested for 5 days and followed by HSV-1 infection, the percentage of viral infected cells were determined by cytometry (a). THP-1 cells were pre-treated with ISRIB, 4μ8C, CaepinA7, then trained with BCG, rested for 5 days and followed by HSV-1 infection (b). THP-1 cells were pre-treated with AMG18 (c) or transfected with siRNA against XBP1 (d), then trained with BCG, rested for 5 days and followed by HSV-1 infection. The level of IFN-β was measured by ELISA (e). THP-1 cells pre-treated with 4μ8C were trained with BCG, then rested for 5 days and followed by VSV infection (f). Data are representative of three independent experiments and showed as mean ± SEM. *p < 0.05, **p < 0.01, ***p < 0.001, ****p < 0.0001 by one-way ANOVA.

In summary, IRE1α-XBP1s pathway of UPR regulates BCG-induced trained immunity against DNA and RNA virus attack by controlling AHR signaling.

Importantly, we observed that IRE1-XBP1s signaling blockade during BCG training in mice impaired control of HSV-infection as determined by the percentage of virally infected cells (Fig. 7a), viral DNA copies (Fig. 7b), viral titers (Fig. 7c) and tissue pathology (Fig. 7d) in lungs. These findings confirmed the role of IRE1-XBP1s pathway on BCG-induced trained immunity against viral infection in vivo.

**Fig. 7.**
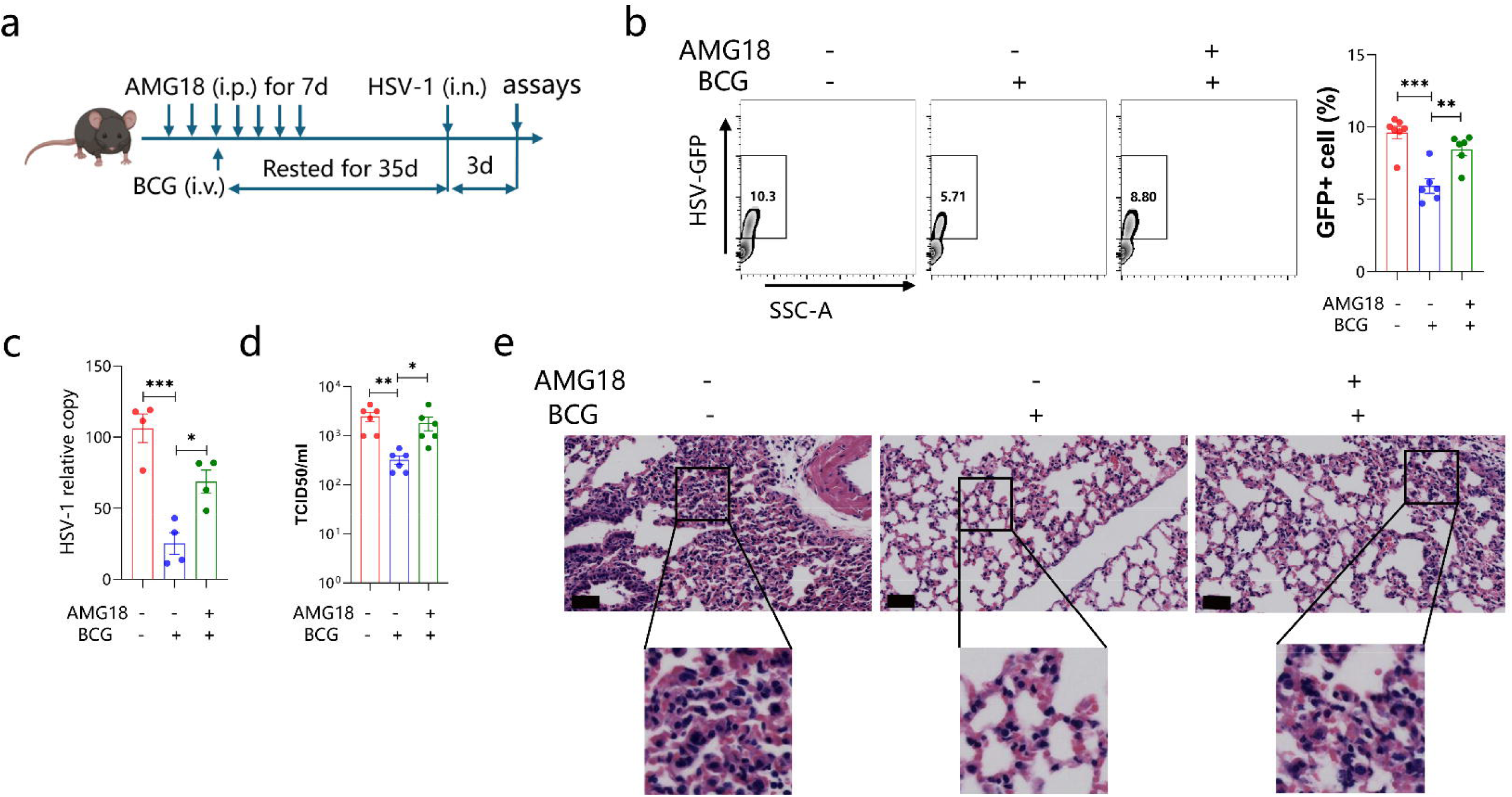
IRE1α-XBP1s signaling regulates trained immunity induced by BCG in vivo. 4-6 mice per group treated with AMG 18 or DMSO as control intraperitoneally every day for serial 7 days were trained with BCG or PBS as control intravenously at the third AMG 18 treatment. Rested for 35 days after BCG training, mice were intranasal infected with HSV-1 for 3 days, lungs were harvested (a) for flow cytometry analysis (b), viral DNA copy assay by quantitative PCR for gD gene (c), viral titers by TCID_50_ assay (d) and H&E staining (e). Scale bar: 50μm. Data are representative of three independent experiments and showed as mean ± SEM. *p < 0.05, **p < 0.01, ***p < 0.001, ****p < 0.0001 by one-way ANOVA.

### Blockade of IRE1α-XBP1s signaling promoted AHR expression and entry into nucleus during BCG training

To explore the mechanism how IRE1α-XBP1s signaling regulates AHR, we assessed the mRNA expression of AHR and its transport protein, aryl hydrocarbon receptor nuclear translocator (ARNT). We observed that antagonizing IRE1α-XBP1s signaling with 4μ8C or AMG18 significantly increased AHR and ANRT mRNA expression at 6 hours or 24 hours after BCG training in vitro (Fig. 8a). Western blot analysis and immunofluorescence assay revealed that BCG training enhanced AHR production and IRE1α-XBP1s inhibition promoted AHR nuclear translocation (Fig. 8b and c and d).

**Fig. 8.**
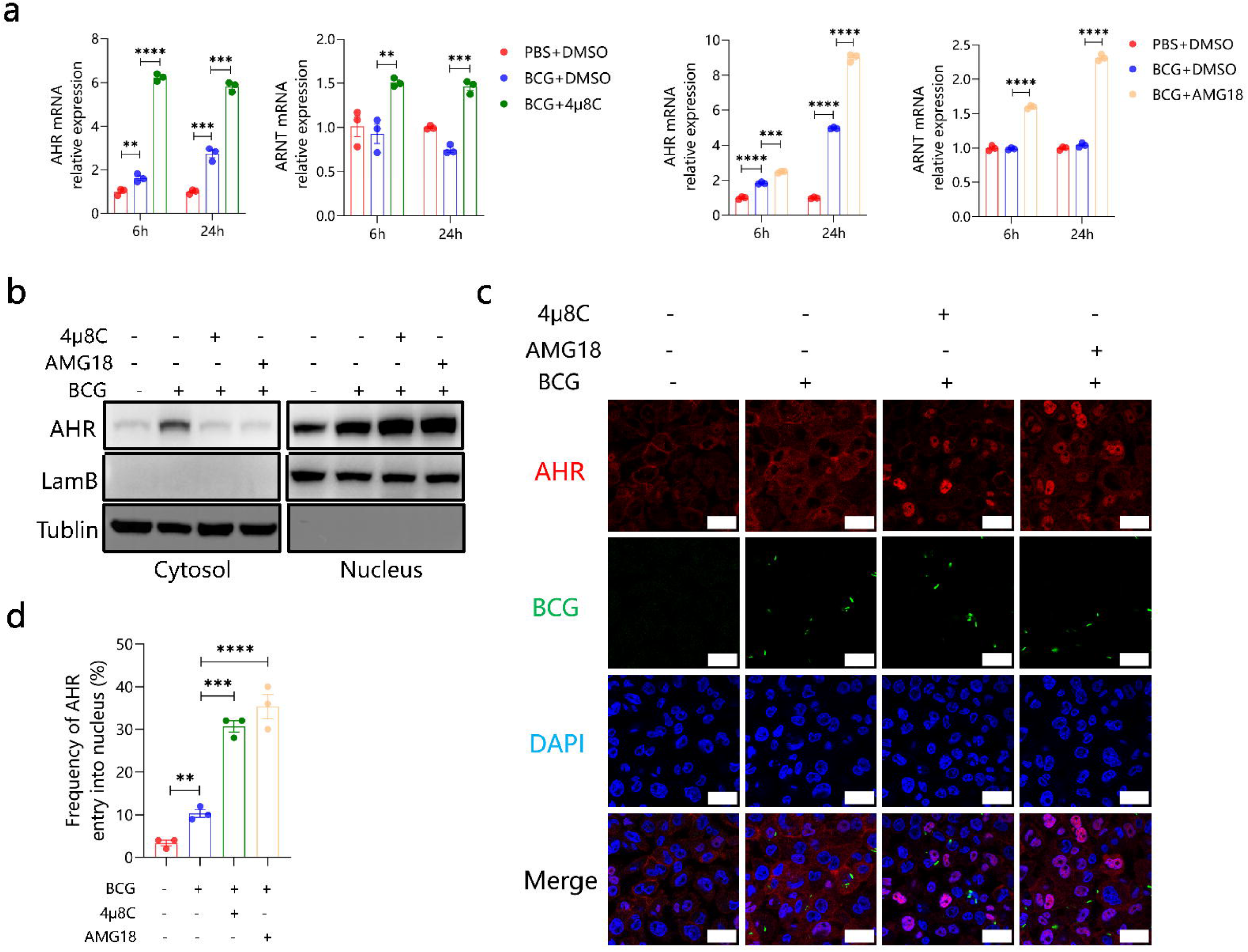
Blockade of IRE1α-XBP1s signaling enhanced AHR expression and entry into nucleus during BCG training. The mRNA expression of AHR or ARNT in THP-1 cells pre-treated with 4μ8C or AMG18 at 6 hours or 24 hours after BCG training were determined by qPCR (a). AHR protein level in cytosol or nucleus were measured by western-bloting (b). Confocal microscopy was used to visualize the location of AHR. Scale bar: 30μm (c). Quantification of the proportion of AHR entry into the nuclei (d). Data are representative of three independent experiments and showed as mean ± SEM. *p < 0.05, **p < 0.01, ***p < 0.001, ****p < 0.0001 by one-way ANOVA.

Thus, IRE1α-XBP1s signaling facilitated BCG-induced trained immunity by negatively regulating AHR through suppressing expression and translocation of AHR into nucleus.

## Discussion

Cells communicate with neighboring cells by paracrine or gap junctions to share information. With GFP-expressing cGAS and mCherry-expressing STING plasmids, cGAMP, which is synthesized by cGAS, could transfer from producing cells to bystander cells through gap junction to active bystanders’ STING and promote their antiviral immunity [12]. Then cGAMP was proved to be able to transfer to bystanders by a volume-regulated anion channel called LLRC8 [13]. In respiratory syncytial virus (RSV) and influenza virus co-infection experiment, RSV-infected cells secreted IFN-β and IFN-λ to protect bystander cells against influenza virus [14]. With a cell line that can secrete mCherry and a GFP-expressing cytomegalovirus (CMV), viral-infected cells had been found to promote the susceptibility of neighboring cells to subsequent CMV, HSV-1 and influenza virus infection by inhibiting neighboring cells’ mitosis and innate immune response and changing the extracellular matrix. Meanwhile, distal cells were better able to slow viral spread due to enhanced antiviral response [15]. Someone has reported that SARS-CoV-2 infected cells could enhance the level of inositol requiring enzyme 1 alpha (IRE1α) dependent NUAK2 of bystander cells by paracrine and then promote ACE2 receptor expression and facilitate susceptibility to SARS-CoV-2 [16]. Nevertheless, the profile of bystander cells in trained immunity remains elusive. In our study, we found that BCG training could protect not only BCG harboring cells but bystander cells against DNA virus (HSV-1) and RNA virus (VSV) infection. At the same time, we also observed an interesting phenomenon that the percentage of viral infection in BCG haboring cells was higher than that of bystander cells upon HSV-1 infection, while the result was inverse on VSV infection. We further provided evidence that this odd phenomenon was due to the fact that BCG training induced increased HSV-1 entry receptors (PILRA, TNFRSF14) expression in BCG harboring cells (Fig. 2, Fig. 9a). Moreover, our results also suggested that bystander cells had no immune memory upon BCG training although they could strongly defend against viral infection compared to the control group (no training) and they acquired protection from BCG harboring cells during viral infection partly via type I IFN signaling. However, other mechanisms independent of type I IFN might be involved in the antiviral capacity induced by BCG training (Fig. 3c, Fig. 4d), which had not yet been further explored in this study.

**Fig. 9.**
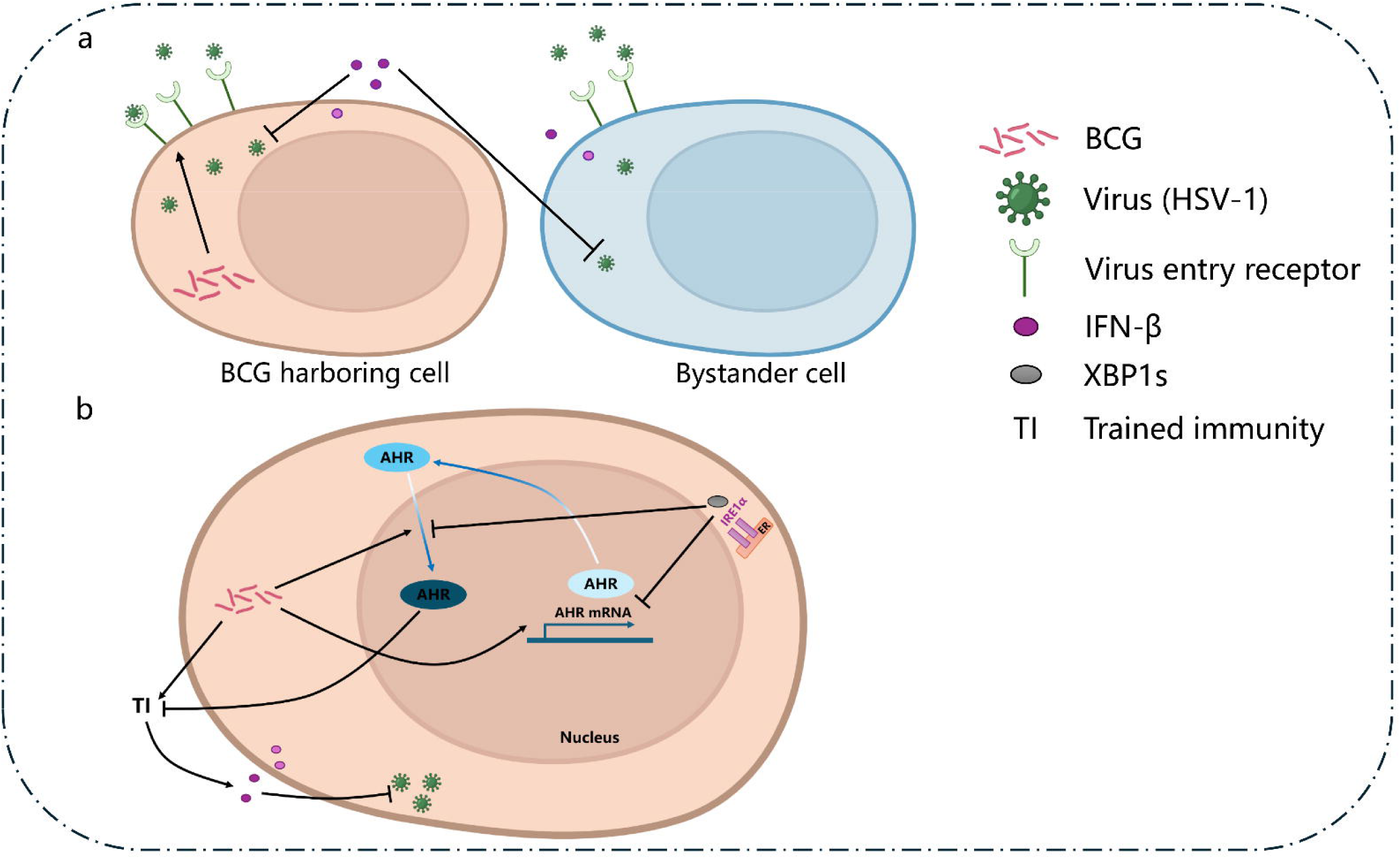
Model depicting the effect of BCG training on BCG harboring cells and bystanders and the role of IRE1α-XBP1s pathway on regulating BCG-induced trained immunity against viral infection. Cells infected with BCG (BCG harboring cells), on the one hand, promotes type I IFN production upon HSV-1 infection, which protects both themselves and bystander cells, on the other hand, enhances HSV-1 entry receptors expression, which leads to an increase in their own viral infection rate (a). IRE1α-XBP1s signaling inhibits AHR mRNA expression and entry into nucleus, which reduces the suppression of AHR signaling on the trained immunity induced by BCG, leading to enhanced production of type I IFN and lower viral load upon virus infection.

Metabolic and epigenetic reprogramming were reported to be two primary molecular mechanisms in the induction of trained immunity. It was reported that glycolysis, oxidative phosphorylation, cholesterol biosynthesis and linoleic acid metabolism were involved in trained immunity [17–21]. These metabolic alterations influenced trained immunity might be through affecting energy providing or substance synthesis involved in immune response or epigenetic modification. With Assay for Transposase Accessible Chromatin sequencing (ATAC-seq) or Chromatin Immunoprecipitation sequencing (CHIP-seq), a good many studies showed that immune stimulus (like BCG, β-glucan or Candida ablbicans et.al.) elicited higher signals or histone modification in the genes associated with proinflammatory response, chemokines [7, 8, 10, 11, 21]. H3K4 me3, H3K27ac were reported to be two primary histone modification in BCG-induced trained immunity [7, 8, 10, 11], while in our study, we found that inhibition of these two epigenetic modifications with selective antagonists could not significantly affect the effect of BCG training upon HSV-1 infection. AHR is a ligand-activated transcript factor that plays an important role in regulating immunity and cell differentiation. In our and other previous studies, AHR was reported to be a negative regulator in immune response in viral infection. Activating AHR signaling downregulated type I IFN-mediated antiviral response [22] and AHR deficiency reduced viral load [9]. While the role of AHR in BCG-induced trained immunity has not been fully elucidated. Here we observed that blockade AHR signaling with selective inhibitor or genetic knockout during BCG training significantly reduced viral infection in vitro and in vivo. This enhanced BCG-induced trained immunity against virus by AHR deficiency partially depended on type I IFN. These results suggested that AHR could be a negative regulator in BCG-induced trained immunity. Furthermore, we found that AHR signaling was regulated by IRE1α-XBP1 signaling (one of the pathway in the unfolded protein response (UPR)) in BCG training. UPR, consisting of three sensors, IRE1α, protein kinase R-like ER kinase (PERK), activating transcription factor 6 (ATF6), is important in response to endoplasmic reticulum (ER) stress and has also been reported to be involved in immune response and diseases [23–26]. IRE1α signaling is the most conserved pathway in UPR. When ER stress happened, IRE1α’s endoribonuclease is activated and then cleaves transcription factor X-box binding protein 1 (XBP1) mRNA, yielding a spliced and active mRNA called XBP1s, XBP1s translocate into the nucleus to active the expression of several genes related to protein folding and immune response [27]. However, the role of UPR signaling in trained immunity has not yet been understood. Here, we demonstrated that IRE1α-XBP1s was critical in BCG-induced trained immunity in vitro and our murine experiment result whereafter confirmed this finding.

Inhibition of IRE1α-XBP1s with antagonist or siRNA during BCG training resulted in increased viral infection and reduced IFN-β level. Our results further suggested that BCG training increased the expression of AHR and suppression of IRE1α-XBP1s signaling impaired BCG-induced trained immunity by enhancing AHR and ARNT mRNA expression and facilitating AHR protein entry into nucleus (Fig. 8, Fig. 9b). Previous studies reported that in macrophages XBP1s could be activated by TLR2 or TLR4 signaling and promote IL-6, TNF, and IFN-β production by recruited to the promoters of IL-6, TNF and IFN-β [28–30]. It was the first time that uncover the mechanism by which IRE1α-XBP1s signaling regulating BCG-induced trained immunity upon viral infection by affecting AHR expression and entry into nucleus. Microorganisms could hijack host’s factors to suppress immune response for their infection and survival [33–35]. Given that BCG which was used as immune stimulus here for inducing trained immunity was also a microorganism, we have reason to speculate that BCG might take advantage of some cellular contents, such as AHR, to evade antimicrobial immunity and impair the intensity of its inducing trained immunity as a consequence. These imply that host factors should be taken into account when conducting trained immunity.

In summary, our study revealed that BCG training could induced trained immunity in macrophages against DNA and RNA virus infection. BCG harboring cells protected bystander cells upon viral infection partly via type I IFN although BCG training could enhance some viral entry receptors expression which resulted in increased viral infection rate in BCG harboring cells. AHR was a negative regulator in BCG training and IRE1α-XBP1s pathway was important in BCG-induced trained immunity by inhibiting of AHR mRNA expression and AHR entry into nucleus.

## Materials and methods

### Cell culture and in vitro trained immunity experiments

THP-1 cells (RRID: CVCL_0006; ATCC® TIB-202^TM^) or U937 cells (RRID: CVCL_0007; ATCC® CRL-1593.2^TM^) were cultured in RPMI 1640 medium supplemented with 10% fetal bovine serum (FBS) and 0.05 mM 2-mercaptoethanol. RAW264.7 cells (RRID: CVCL_0493; ATCC® TIB-71^TM^) or A549 cells (RRID: CVCL_0023; ATCC® CCL-185^TM^) were cultured in DMEM medium supplemented with 10% FBS. In vitro trained immunity experiments, cells were activated with 100 nM phorbol 12-myristate 13-acetate (PMA) (P1585, Sigma-Aldrich) for 24 hours to differentiate into macrophages and rested for 24 hours. Then the macrophages were trained with *M. bovis* BCG Pasteur for 24 hours and washed and rested for 5 days for further viral infection.

### Chemicals and antibodies

CH223191 (S7711, Selleck), ISRIB (HY-12495, MCE), 4μ8C (SML0949, Sigma-Aldrich), CaepinA7 (HY-108434, MCE), AMG18 (HY-114368, MCE) or APY29 (HY-17537, MCE) were used 1 hours prior to BCG training at indicated concentration and present for another 24 hours during the training period.

Anti-IFNAR antibody (RRID: AB_387828; 21385-1, PBL assay science) for block type I IFN signaling was used at HSV-1 infection. Anti-phospho-IRF3 antibody (RRID: AB_2773013; 29047, Cell Signaling Technology), anti-IFN-β (RRID: AB_2799843; 73671, Cell Signaling Technology), anti-PILRA antibody (RRID: AB_2576622; PA5-42954, thermo), anti-TNFRSF14 (RRID: AB_2866601; MA5-35989, thermo), anti-LDLR antibody (RRID: AB_2809369; MA5-32075, thermo) and anti-rabbit IgG (H+L)-Alexa Fluor 647 (RRID: AB_10693544; 4414, Cell Signaling Technology) were used for cytometry analysis.

Anti-AHR antibody (RRID: AB_2800011; 83200, Cell Signaling Technology), anti-LAMB1 antibody (RRID: AB_3698147; 83912, Cell Signaling Technology), anti-α-Tubulin (RRID: AB_2619646; 2125, Cell Signaling Technology) were used for immunoblotting or immunofluorescence.

### Immunofluorescence and flow cytometry

For immunofluorescence staining, after being fixed with 4% paraformaldehyde (PFA) for 15 minutes, the cells were permeabilized with Intracellular Staining permeabilization Wash Buffer (421002, biolegend) for 1 hour and then inoculated with anti-AHR antibody for 2 hours, then stained with the second antibody anti-rabbit IgG (H+L)-Alexa Fluor 647 and 4,6-diamidino-2-phenylindole (DAPI) for 1 hours. Imaging was performed with confocal microscope (LSM980, Zess).

For flow cytometry, generally, cells were fixed with 4% PFA for 15 minutes and washed with phosphate-buffered saline (PBS), resuspended in PBS, and analyzed by cytometry (FACSymphony A3, BD). Intercellular staining was performed according to a procedure described in previous reports [31, 32].

### Virus titer determination

The virus titter was determined by TCID_50_ assay in Vero cells. Briefly, cells were infected with HSV-1 or VSV for certain times, harvested and lysed. The lysates were 10-fold serially diluted and transferred into Vero cells and cultured at 37℃ in a humidified atmosphere with 5% CO_2_ and checked daily until cytopathic effect (CPE) had no more progress. The viral titter was calculated with the Reed-Muench method.

### Enzyme-linked immunosorbent assay (ELISA)

The level of IFN-β in culture supernatants was performed by commercial ELISA kit (KE00195-96T, Proteintech), and assayed using a microplate reader.

### Quantitative real-time PCR

Total RNA was extracted using a total RNA kit (R6834, Omega) according to the manufacturer’ s instructions, followed by cDNA synthesis and real-time PCR assay with SYBR reagent on instrument (ABI7500, thermo). All the primer sequences are available in the supplemental material.

### Immunoblotting

Cells were harvested and protein from cytosol or nucleus were extracted according to manufacturer’ s instructions (P0028, Beyotime). The samples were detected by Western blotting with primary antibodies against AHR, LAMB1, α-Tubulin for overnight and then with secondary antibody against rabbit IgG for 1 hours. The proteins were detected by using a chemiluminescent substrate.

### Animal experiment

All mice were housed in specific pathogen-free conditions, and our experimental procedures were approved by the animal ethics committee of the Third People’s Hospital of Shenzhen (the animal experiment ethics number: 2022-084). Mice aged between 8 to 10 weeks were trained with a single dose of 10^6^ colony-forming units (CFU) of BCG per mouse intravenously. After rested for 35 days, mice were infected with 10^8^ plaque-forming units (PFU) of HSV-1 intranasally for 3 days and lungs were harvested for further assays.

### Statistical analyses

Data are shown as means ± standard errors of the means (SEM), and P values of less than 0.05 were considered significant. Significance was determined by two-tailed Student’s t test unless otherwise stated. GraphPad Prism 6 software (GraphPad Software, CA, USA) was used for data analysis.

## Supporting information

primers-sequences

## Acknowledgements

We thank Professor Carl G. Feng for providing suggestions and comments to this manuscript. This work was supported by the National Key Research and Development Plan (No. 2021YFA1300902, 2022YFC2304400), the Guangdong Science Fund for Distinguished Young Scholars (No. 0620220214), the Shenzhen Scientific and Technological Foundation (No. KCXFZ20211020163545004, RCJC20221008092726022), the Sanming Project of Medicine in Shenzhen (No. SZZYSM202311009, SZZYSM202311033), the Shenzhen Medical Research Fund (No. B2402040), the Shenzhen Clinical Research Center for Emerging Infectious Diseases (No. LCYSSQ20220823091203007), and the Shenzhen High-level Hospital Construction Fund (SZSM202311033).

## Author contributions

F.W. and G.Z. conceived the project; J.C., C.P., L.Z. and H.X. designed the experiment and performed the experiments and acquired and analyzed data; S.L., J.H., M.H. and Y.Y. assisted with data analysis. J.C. wrote this manuscript. C.P., L.Z. and H.X. revised this manuscript. G.Z. supervised all phases of the project.

