## supplemental Figures for "IRE1α-XBP1s axis is required for BCG-induced trained immunity against viral infection by inhibiting AHR signaling"

**Supplemental Fig.1. Transcriptome characteristics of THP-1 trained with BCG upon HSV-1 infection.** Principal component analysis (PCA) of global genes expression (a). Volcano plot for global genes expression profile (b). Go cellular component analysis between BCG trained THP-1 cells and control group upon HSV-1 infection (c). Kegg_pathway classification analysis between BCG trained THP-1 cells and control group upon HSV-1 infection (d). Heatmap of genes involved in immune system and infectious disease (viral) (e).

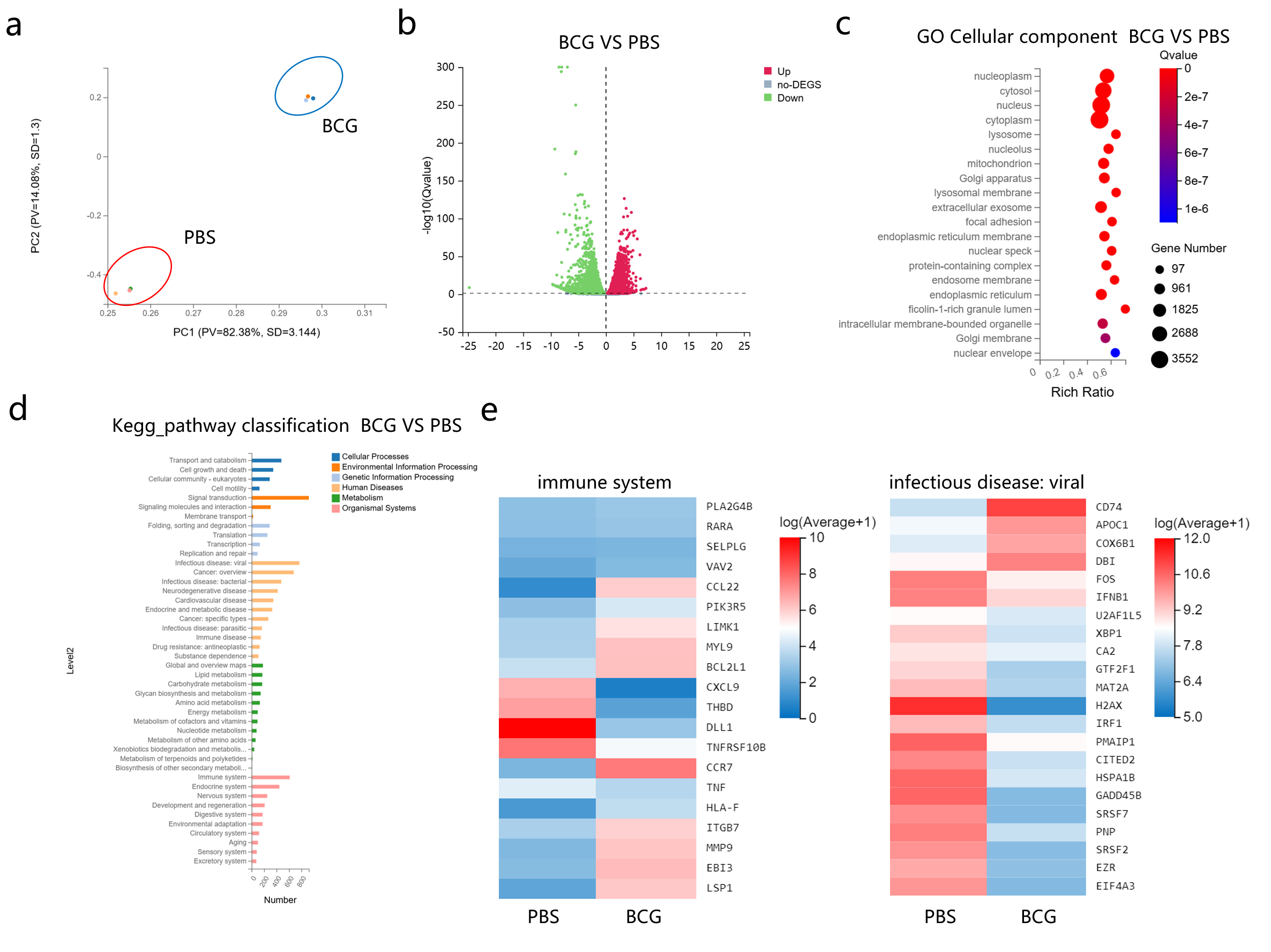

**Supplemental Fig.2. BCG harboring cells express higher HSV-1 receptors that result in increased HSV-1 infection.** Percentage of IFN-β positive cells in BCG harboring cells and bystander cells in BCG trained group upon HSV-1 (a) or VSV infection (b). Heatmap for the expression of HSV-1 and VSV receptors (c). The percentage of PILRA positive cells, TNFRSF14 positive cells, LDLR positive cells between BCG harboring cells and bystander cells were compared (d).

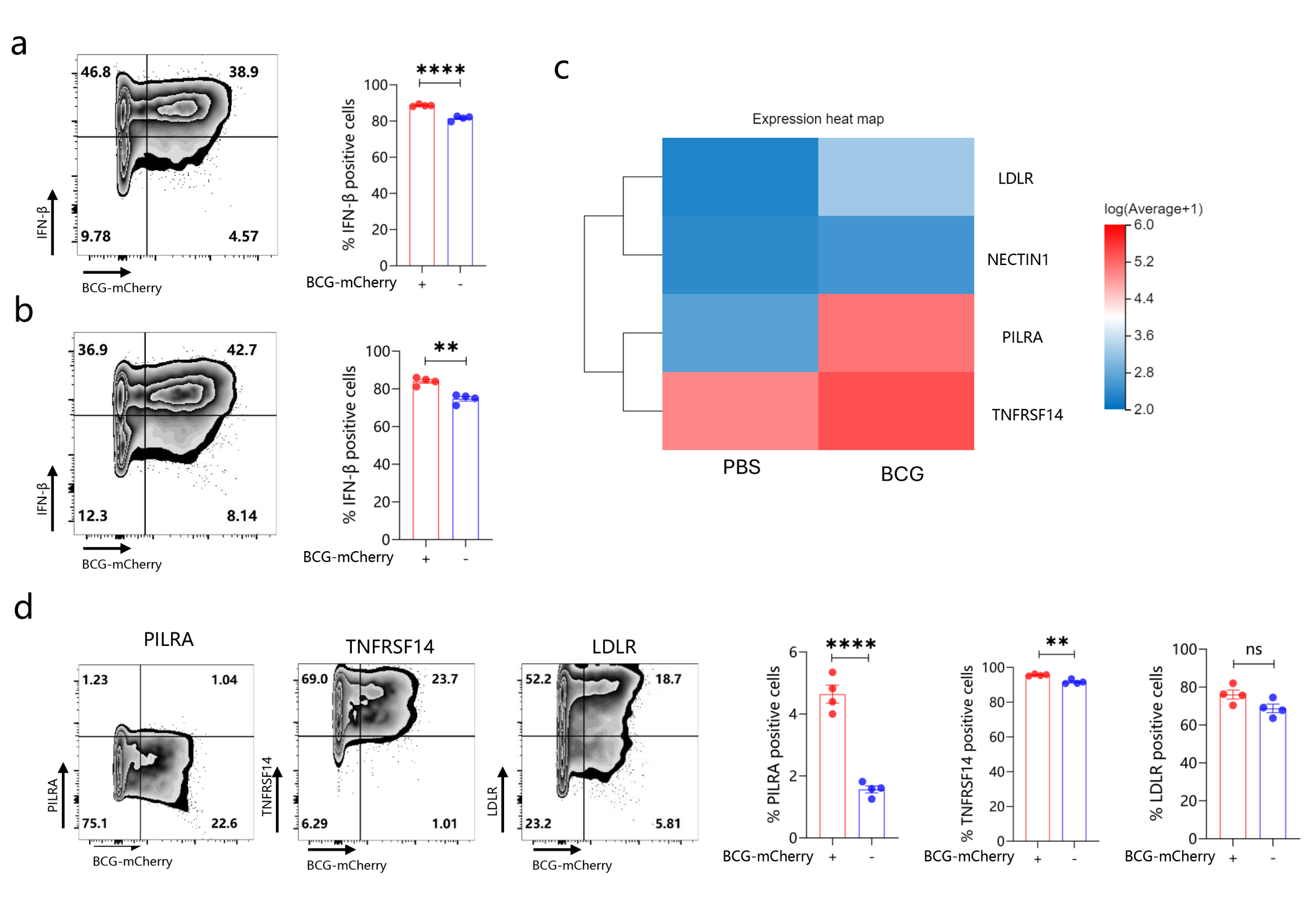

**Supplemental Fig.3. Bystander cells may not have immune memory (trained immunity).** Schematic of the experimental setup (a). HSV-1 viral load between cells inoculated with supernatant from BCG trained cells and control were analyzed by cytometry. THP-1 (b), U937 (c), RAW264.7 (d). VSV viral load between cells inoculated with supernatant from BCG trained cells and control were analyzed by cytometry in THP-1 (e).

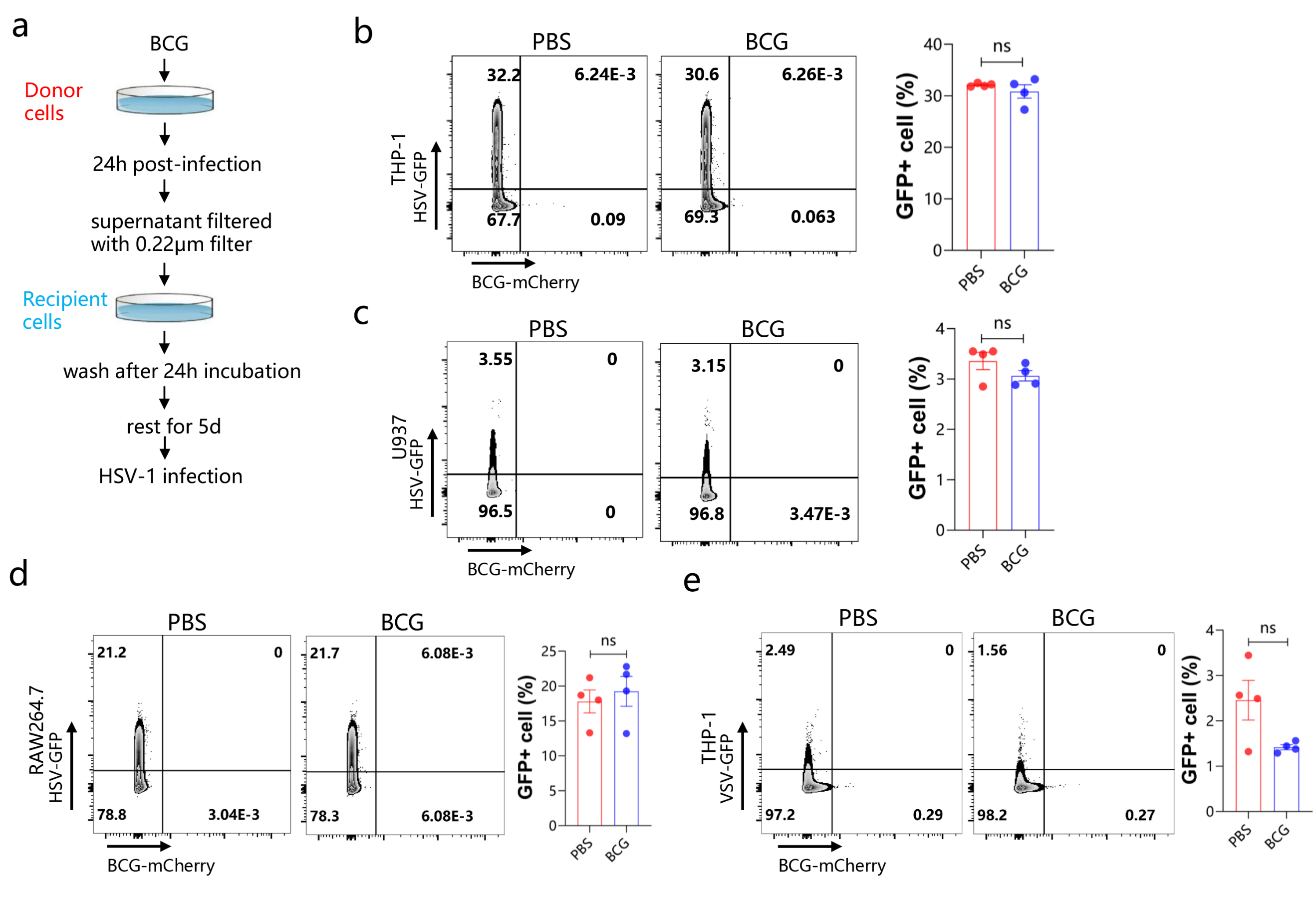

**Supplemental Fig.4. AHR deficiency enhanced BCG induced trained immunity against viral infection might not via altering epigenetics.** THP-1-differentiated macrophages pre-treated with MTA (histone methyltransferases antagonist) or SGC-CBP30 (CBP/p300 bromodomain inhibitor) were trained with BCG for 24 hours and rested for 5 days, followed by HSV-1 infection and viral load (a) and IFN-β level (b) were assayed .

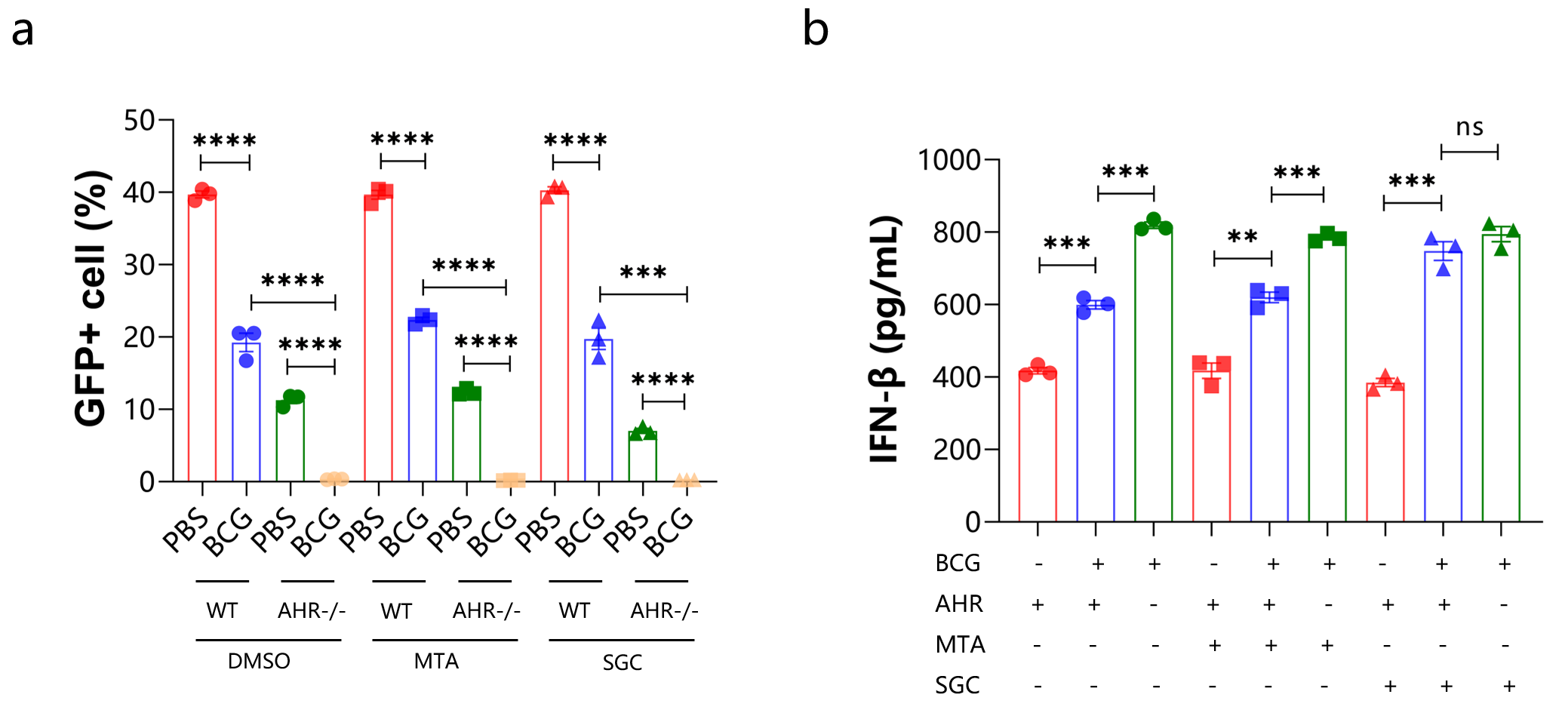
